# FleQ-Dependent Regulation of the Ribonucleotide Reductase Repressor *nrdR* in *Pseudomonas aeruginosa* During Biofilm Growth and Infection

**DOI:** 10.1101/2025.06.03.657667

**Authors:** Domingo Marchan, Alba Rubio, Lucas Pedraz, José María Hernández, Joana Admella, Eduard Torrents

## Abstract

Ribonucleotide reductases (RNRs) are essential enzymes involved in DNA synthesis and repair, catalyzing the conversion of ribonucleotides to deoxyribonucleotides (dNTPs). While all living cells possess at least one RNR encoded in their genome, certain organisms, such as *Pseudomonas aeruginosa,* encode multiple RNR classes. This multiplicity provides a competitive advantage, allowing these organisms to adapt and colonize different environments. Despite their importance, the mechanisms coordinating the expression of different RNRs in microorganisms with multiple RNR classes remain poorly understood. The transcriptional regulator NrdR has been implicated in controlling the expression of all three RNR classes in *P. aeruginosa* by binding to conserved NrdR boxes within the promoter regions of the RNR genes.

To gain insights into the regulation of the different RNR genes, it is first necessary to understand how *nrdR* itself is transcriptionally regulated. In this study, we employed a bioinformatics approach to identify potential transcription factors (TF) involved in *nrdR* regulation. We combined this with promoter-probe vectors *nrdR* promoter fusions to investigate *nrdR* transcriptional regulation and identify TFs that modulate its expression in vitro. Our analysis identified four potential TF that could regulate *nrdR,* and we experimentally confirmed that FleQ is responsible for regulating *nrdR* expression under aerobic and anaerobic conditions. Furthermore, we explored *nrdR* regulation under biofilm-forming conditions and in the *Galleria mellonella* infection model to gain insights into how *nrdR* might be regulated *in vivo*.

**Importance:** This study reveals a nuanced regulatory mechanism by which the transcription factor FleQ, modulated by intracellular c-di-GMP levels, governs the expression of the essential gene *nrdR* in *Pseudomonas aeruginosa*. By demonstrating that FleQ acts as an activator under planktonic and infection conditions and as a repressor during biofilm formation, the findings reveal a dual regulatory role that aligns with the bacterium’s transition between acute and chronic infection states. This dynamic control of *nrdR*, a key repressor of ribonucleotide reductases, links environmental sensing to nucleotide metabolism, offering new insights into how *P. aeruginosa* adapts to diverse and hostile environments. These results not only deepen our understanding of bacterial gene regulation but also highlight potential targets for disrupting biofilm-associated persistence in clinical settings.

## Introduction

*Pseudomonas aeruginosa* is a Gram-negative bacterium of growing medical concern. As an opportunistic pathogen, *P. aeruginosa* is responsible for both acute and chronic infections, including those affecting cystic fibrosis (CF) patients or other forms of bronchiectasis (1, 2). Its remarkable adaptability to diverse environments is largely due to its ability to finely regulate gene expression in response to environmental stimuli (3). However, its prevalence in hospital settings, where antibiotics are frequently used, poses a significant risk, as it rapidly acquires antibiotic resistance (4). The rise of antibiotic-resistant strains has prompted the development of alternative strategies to combat this pathogen (5). One such strategy involves identifying essential molecular targets critical for bacterial survival.

DNA synthesis and repair are vital for all organisms, and the only known biochemical pathway for *de novo* synthesis of deoxyribonucleotide triphosphates (dNTPs) is catalyzed by the enzyme ribonucleotide reductase (RNR) (6). RNRs reduce ribonucleotides to deoxyribonucleotides, the building blocks required for DNA synthesis and repair. Without this enzymatic activity, bacterial viability is compromised. RNRs are divided into three classes (I, II, and III), based on their structural characteristics, metallocofactor requirements, and radical generation mechanisms (6–9). Remarkably, *P. aeruginosa* encodes all three RNR classes (Ia, II, and III) in its genome (10–12). This unique feature provides *P. aeruginosa* with a versatile metabolic toolkit to adapt to fluctuating environments during infection, making RNRs attractive molecular targets.

In recent decades, significant progress has been made in understanding the regulation of RNRs, particularly at the transcription level. Several regulators have been identified that modulate the expression of *nrd* genes. Among these is NrdR, a negative transcriptional regulator of RNR expression (6, 13, 14).

NrdR was first identified upstream of the class II RNR gene (*nrdJ*) in *Streptomyces coelicolor* and *Streptomyces clavuligerus* (15). It is broadly conserved across the bacterial domain, and its genomic location is often found clustered with the *nrd* genes, near DNA replication genes (*dnaB* and *dnaI*), or adjacent to riboflavin biosynthetic operons, depending on the species (14, 16). NrdR regulation depends on specific DNA motifs called NrdR-boxes (14), which are typically present in tandem, within promoter regions, although some genes may be regulated by a single box.

The NrdR protein, composed of 140-200 amino acids, has a well-characterized structure (17). It contains two main domains: an N-terminal Zn-finger domain responsible for DNA binding and a central ATP-cone domain that resembles the allosteric activity domain of RNRs (6, 17, 18). The ATP-cone domain plays a regulatory role linked to intracellular dNTP concentrations (11, 17). Conformational changes in this domain, triggered by fluctuations in dNTP levels, modulate DNA binding activity. Studies have shown that NrdR regulation involves the cooperative and allosteric binding of ATP (adenosine triphosphate) and dATP (deoxyadenosine triphosphate) (18). Additionally, the ATP-cone domain mediates NrdR oligomerization, which directly influences its DNA-binding capacity (17).

Despite these advances, many aspects of *nrdR* regulation remain poorly understood across bacterial species. For example, our recent work demonstrated that *nrdR* expression in *P. aeruginosa* is negatively regulated under anaerobic conditions by the transcriptional factor NarL (11).

In this study, we combined a bioinformatics-based transcription factor screening with in vitro binding assays to identify regulators that directly interact with the *nrdR* promoter. Our findings reveal that FleQ, a previously known regulator of motility and biofilm formation, also modulates *nrdR* expression. We further investigated the role of FleQ-mediated *nrdR* regulation under biofilm growth conditions and validated its functional significance *in vivo* using the *Galleria mellonella* infection model.

## Results

### *nrdR* gene context overview and bioinformatic identification of transcription factors

This study aims to elucidate the transcriptional regulation of the *nrdR* gene and to identify potential transcription factors involved in modulating its expression. The *nrdR* gene (PA4057) is part of the *nrdR*-*ribD* operon, as previously described (Figures 1A–B) (11). To precisely define the *nrdR* promoter region, we first determined its transcriptional start site (TSS) experimentally using the 5′- RACE technique (see Materials and Methods). The TSS (+1) was mapped to 28 bp upstream of the translational start codon (Figure 1D and Supplementary Figure S1) under both aerobic (91.7% of sequences) and anaerobic (50% of sequences) conditions. These results corroborate previously published data obtained under the same oxygenic conditions (28).

**Fig 1.**
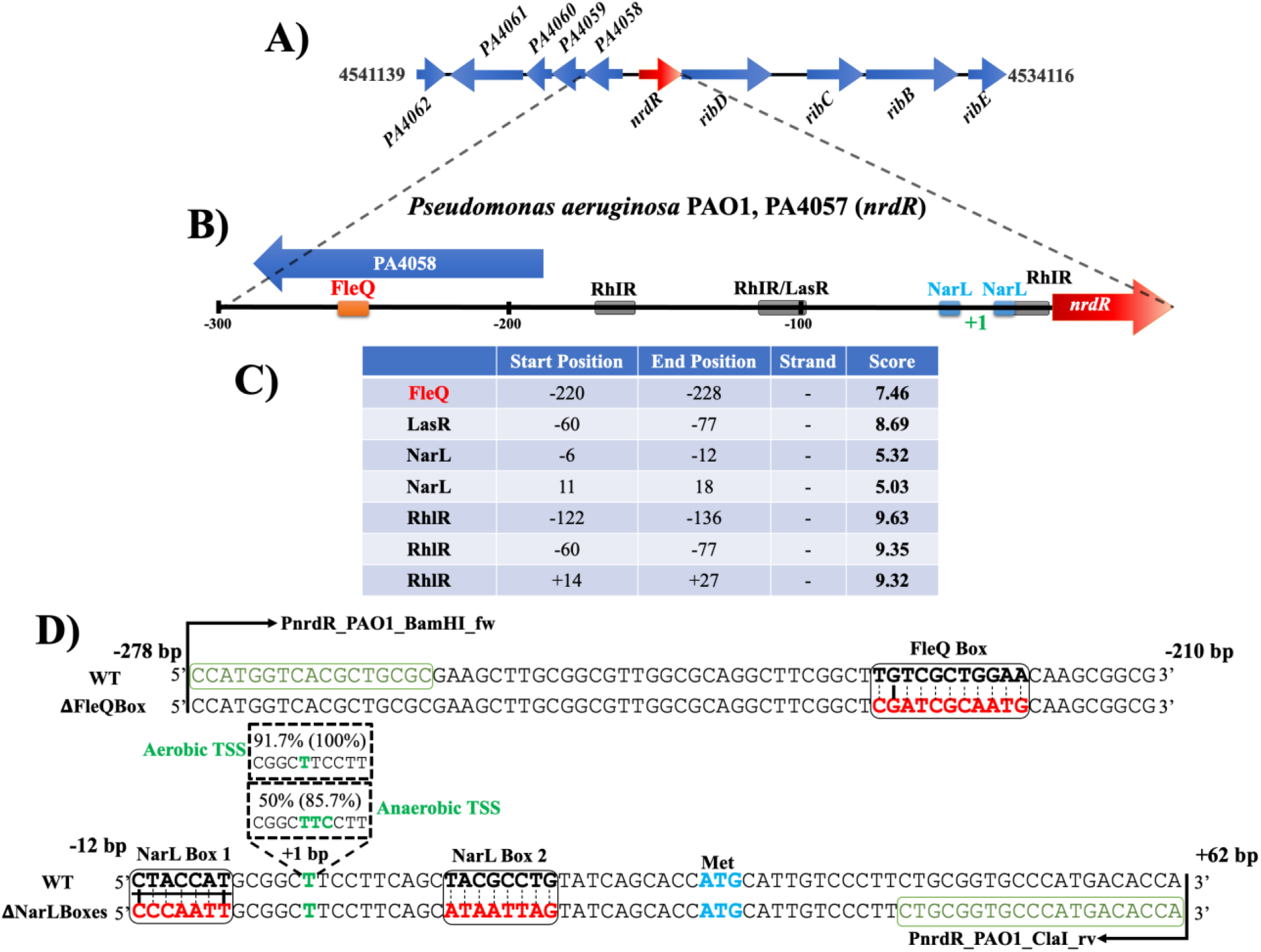
*nrdR* operon organization and transcription factor binding site predictions. A) Schematic of the *nrdR-ribD* operon^12^. B) Predicted transcription factor binding sites in the *nrdR* promoter región. C) Virtual Footprint analysis using PRODORIC2 with *P. aeruginosa* weight matrices (PWM). D) Detailed promoter map showing predicted FleQ and NarL binding sites (black bold), mutated sequences (red), -10 box (underlined), ATG start codon (bold blue), and TSS (+1) (green) identified under aerobic or anaerobic conditions.

Following TSS mapping, we performed *in silico* analysis using the PRODORIC2 Virtual Footprint tool to identify putative transcription factor binding sites in the promoter region. We identified seven binding motifs: one undecameric FleQ-binding site (-224 bp upstream of the TSS, referred to as the FleQ-box TGTCGCTGGAA); one heptadecameric LasR-binding site (-68 bp upstream, LasR-Box CTCTCGCCTTTCCCAGC); three RhlR-binding sites (-129, -68 and +23 bp upstream, RhlR-Boxes CTTGTATAGCGCAA, CTCTCGCCTTTCCCAGC, CTGTATCAGCACC); and two heptameric NarL-binding sites previously described (11) (Figure 1B-C). The role of FleQ will be explored further in this study.

The two NarL boxes are located near the predicted -10 region (CTACCAT; Figure 1D), and we have previously described their implication in anaerobic activation of *nrdR* transcription (11). To evaluate the potential involvement of quorum sensing (QS) regulators LasR and RhlR, we assessed *nrdR* expression in a *P. aeruginosa* PAO1 Δ*lasI/rhlI* mutant. No significant expression differences were observed between wild-type and the QS-deficient mutant (Supplementary Figure S2). Given the prevalence of false positives in motif prediction, we ruled out any role for quorum sensing in *nrdR* regulation based on our experimental data.

### The pull-down confirms binding of FleQ and NarL to the *nrdR* promoter

To confirm direct binding of FleQ to the *nrdR* promoter, we initially attempted an electrophoretic mobility shift assay (EMSA). However, FleQ overproduction consistently yielded insoluble protein aggregates (inclusion bodies) under various overproduction conditions (Supplementary Figure S3). As an alternative, we performed a protein pull-down assay coupled with LC-MS proteomic analysis to identify *in vivo* DNA-binding proteins interacting with the *nrdR* promoter.

We used a biotinylated DNA fragment containing three tandem copies of the P*nrdR* sequence (from -306 pb to +34 pb) immobilized on streptavidin-coated magnetic beads (see Materials and Methods and a summary in Supplementary Figure S4). Protein extracts from *P. aeruginosa* PAO1 cultures grown to exponential and stationary phases were incubated with the bait DNA. Six different bands were resolved via SDS-PAGE from stationary samples, corresponding to 50 kDa (Band 1), 37 kDa (Band 2), 27 kDa (Band 3), 24 kDa (Band 4), 15 kDa (Band 5), and 13 kDa (Band 6) (Figure 2A).

**Fig 2.**
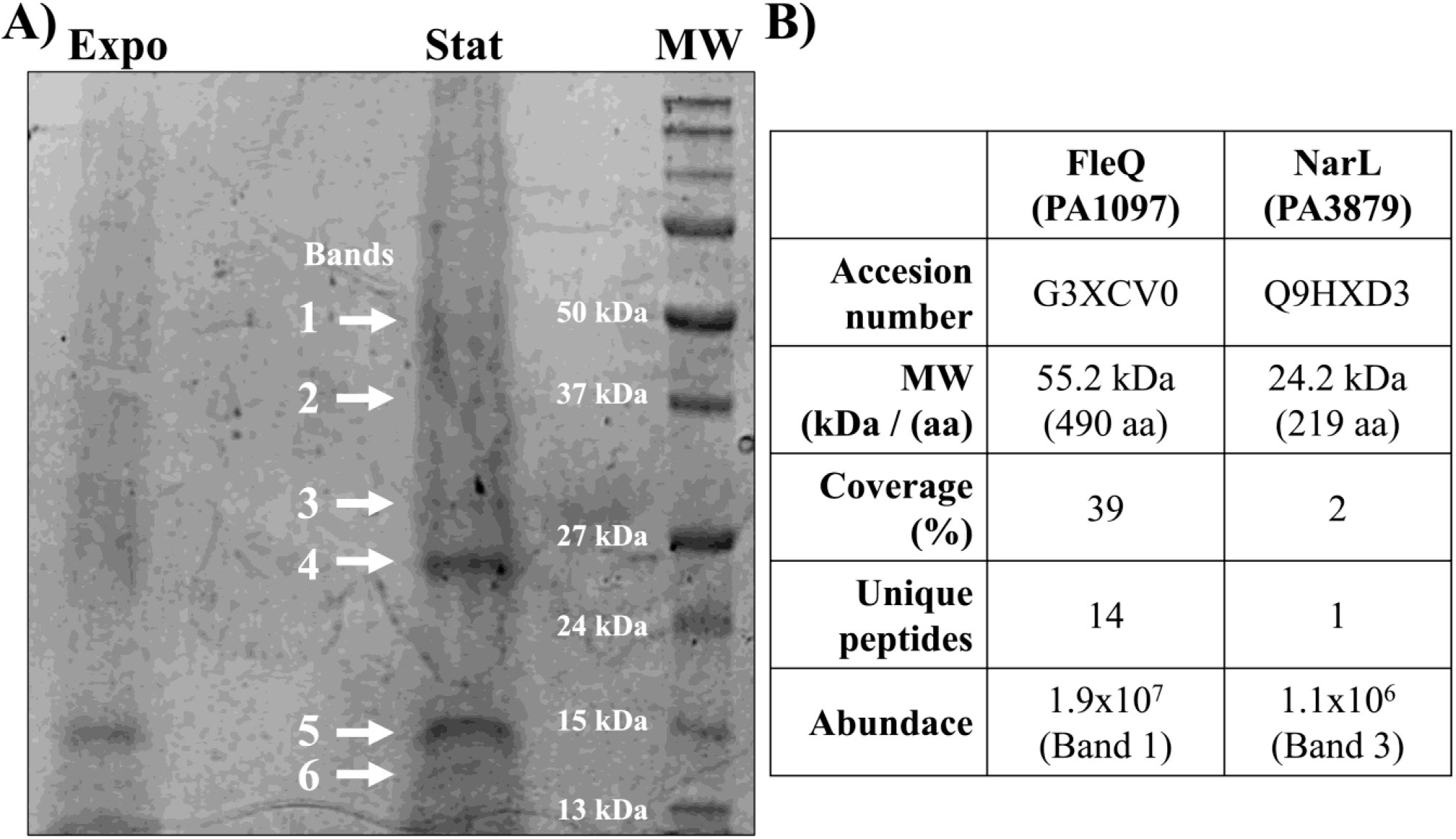
SDS-Page gel from *nrdR* promoter pull-down assay. A) Protein extracts from *P. aeruginosa* PAO1 grown in exponential (Expo) and stationary (Stat) phases were incubated with a biotinylated *nrdR* promoter. After the pull-down, the protein extract was resolved on a 12% SDS-Page gel. Black arrows indicate the excised bands analysed. B) Summary of the LC-MS hits regarding the FleQ and NarL. Molecular weight (MW), peptide coverage (%, indicating the percentage of peptides detected), and abundance (m/z) (total amount of protein detected in each band) values are provided.

Proteins were identified via LC-MS using the *P. aeruginosa* PAO1 (UP000002438) proteome as a reference (see Materials and Methods). A Supplementary Table III with complete proteomic data is available as supplementary material. FleQ was detected with 14 unique peptides identified, covering 39% of the entire protein in Bands 1–3, with the majority in Band 1, corresponding to its apparent molecular weight (Supplementary Figure S5). NarL was also identified in Band 3 with 5% coverage, although not at its optimal molecular weight, which is why this low-abundance identification is shown in Figure 2B. These results confirm the binding of FleQ and NarL to the *nrdR* promoter region. Notably, LasR and RhlR were not identified in any band with sufficient coverage, supporting the hypothesis that they do not bind this region under the tested conditions.

### FleQ activates *nrdR* transcription under aerobic and anaerobic conditions

To investigate the functional impact of FleQ on *nrdR* transcription, we monitored bacterial growth in *P. aeruginosa* PAO1 and its Δ*fleQ* mutant (PW2981) under aerobic and anaerobic conditions (Figure 3A). No significant differences in growth were observed under aerobic conditions. However, under anaerobic conditions, the Δ*fleQ* strain exhibited delayed entry into the stationary phase, with a growth lag of ∼3 hours compared to wild-type. These findings should be taken into account when interpreting the results of subsequent experiments conducted under these specific conditions.

**Fig 3.**
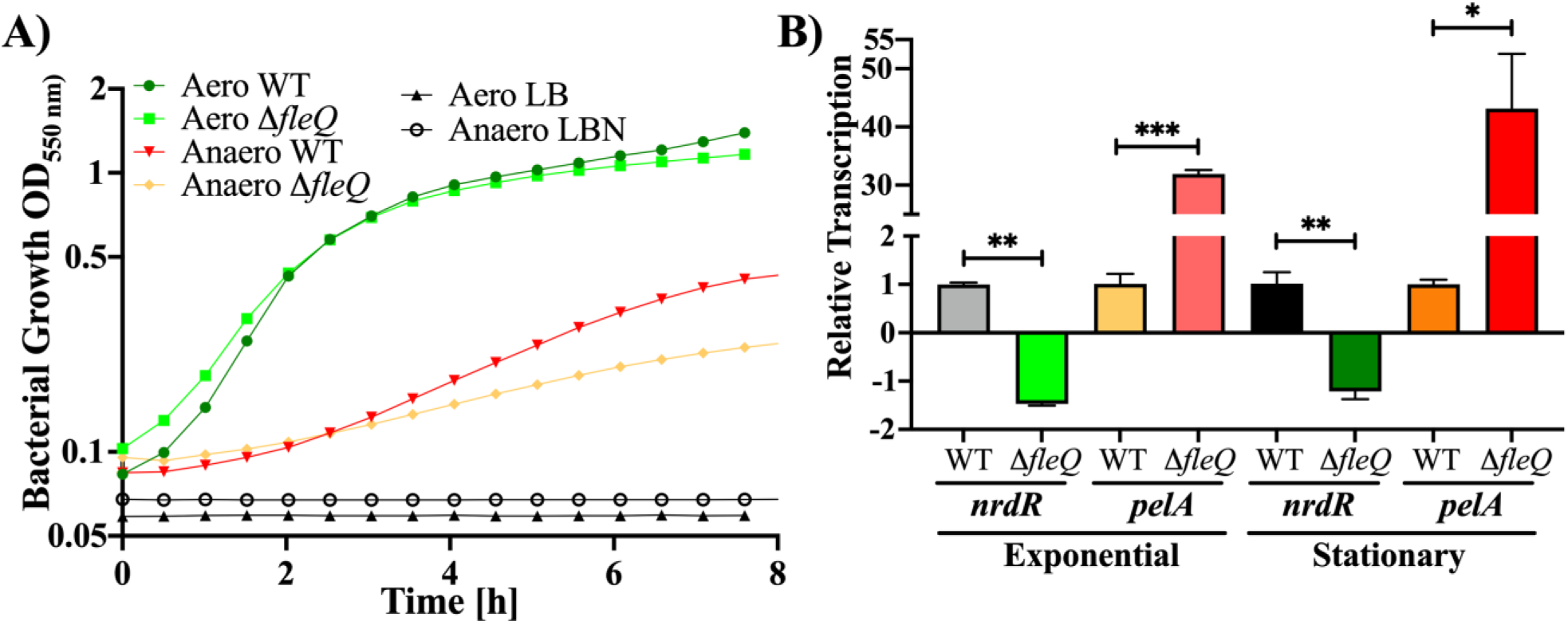
Growth and transcriptional analysis of *nrdR*. A) Growth curves of *P. aeruginosa* PAO1 wild-type and isogenic Δ*fleQ* strains under aerobic and anaerobic conditions. Negative controls were LB or LBN without cells. B) RT-qPCR of *nrdR* and *pelA* in both *P. aeruginosa* PAO1 Δ*fleQ* vs wild-type strains under exponential (OD_550_= 0.45) and stationary (OD_550_= 2.5) aerobic growth phase. Expression values were normalized to *gapA.* Error bars represent standard deviation from three independent biological replicates. Statistical analysis was performed using Student’s unpaired *t*-test (*, *p* < 0.05; **, *p* < 0.01; ***, *p* < 0.001).

We then measured *nrdR* mRNA levels in both *P. aeruginosa* PAO1 wild-type and its isogenic Δ*fleQ* mutant by RT-qPCR, using *pelA* (a known FleQ-regulated gene) as a positive control and *gapA* as an internal control (see Materials and Methods). Samples were collected aerobically during the exponential (OD₅₅₀ = 0.45) and stationary (OD₅₅₀ = 2.5) phases, corresponding to active and inactive celullar DNA replication, respectively, and reflecting active or inactive RNR enzyme activity. The *pelA* gene, which produces the exopolysaccharide required for biofilm formation, contains two experimentally identified FleQ binding sites in its promoter region. As expected, *pelA* was significantly upregulated (30-45 times) in the Δ*fleQ* mutant, consistent with FleQ’s repressor role during planktonic growth (29, 30). In contrast, *nrdR* expression was moderately reduced in the Δ*fleQ* mutant under both growth conditions, indicating that FleQ acts as a positive regulator of *nrdR* (Figure 3B).

We next generated *PnrdR*–GFPmut3 transcriptional fusions on plasmid pETS130 (pETS130:P*nrdR*(PAO1)) to generate fluorescent-based reporter assays (see Material and Methods). A version with a mutated FleQ binding site (pETS130:*PnrdR*ΔFleQBox) was also constructed (see Materials and Methods). Reporter assays confirmed that deletion of *fleQ* or mutation of the FleQ-box reduced promoter activity under both aerobic and anaerobic conditions (Figure 4A–B). FleQ complementation (pUCP20T:P*fleQ*:*fleQ*) restored expression levels. These results reinforce that FleQ directly activates *nrdR* transcription as also observed in the RT-PCR (Figures 3A and 4A).

**Fig 4.**
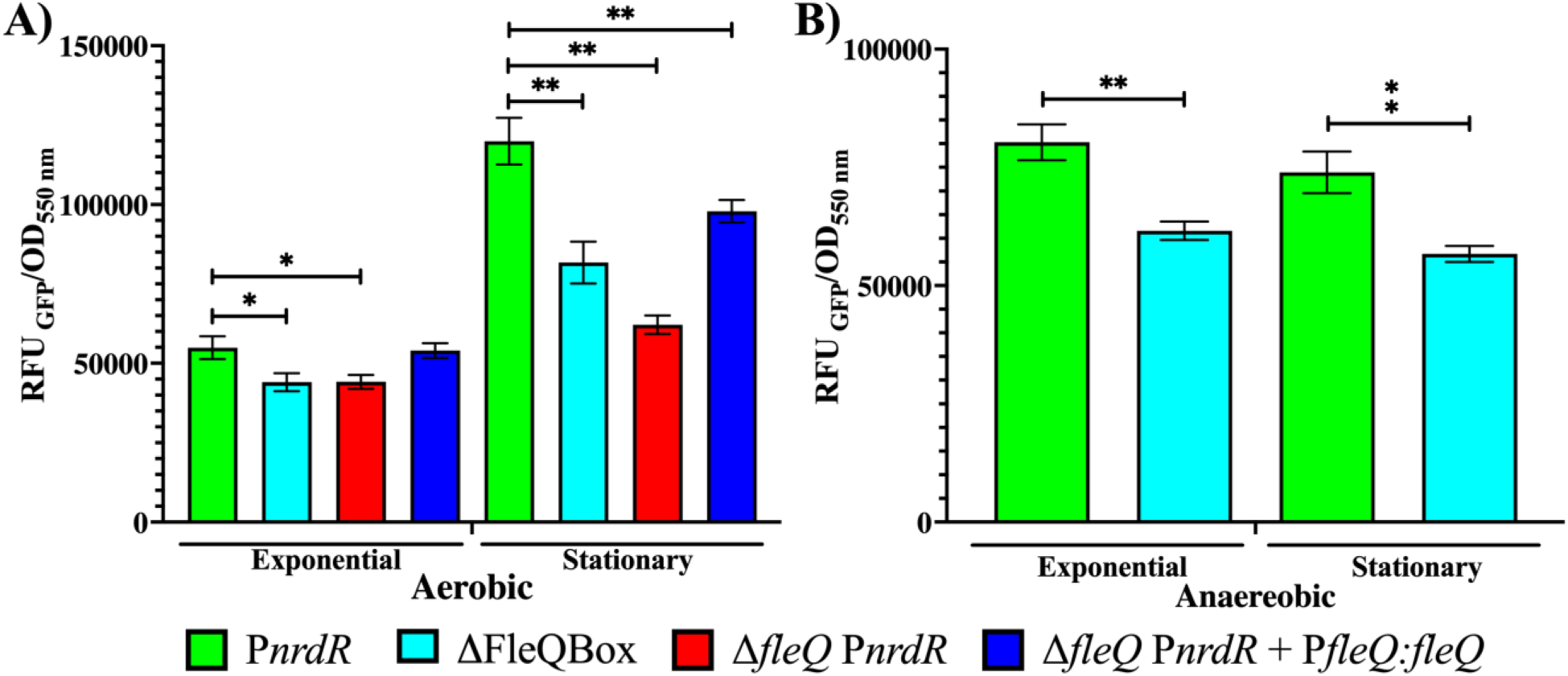
*nrdR* promoter activity measured by GFP fluorescence. A) Aerobic conditions in exponential (OD_550_ 0.45) and stationary (OD_550_ >2.5) of transcriptional fusions between P*nrdR* promoter [pETS130:P*nrdR*(PAO1), green] and P*nrdR* ΔFleQBox [pETS130:P*nrdR*(PAO1) ΔFleQBox, cian blue]. The *P. aeruginosa* PAO1 Δ*fleQ* strain [pETS130:P*nrdR*(PAO1)] and its complemented strain [pUCP20T:P*fleq:fleq*] are represented in red and dark blue, respectively. B) Anaerobic conditions in exponential (OD_550_ 0.45) and stationary (OD_550_ >2.5) gene reporter assay of transcriptional fusions between P*nrdR* promoter and P*nrdR* ΔFleQBox. Error bars indicate the standard deviation from three independent experiments. Statistical analysis was performed using Student’s unpaired *t*-test (*, *p* < 0.05; **, *p* < 0.01).

### c-di-GMP modulates FleQ-mediated regulation of *nrdR*

FleQ function is tightly regulated by c-di-GMP levels in response to environmental signals (29–31). To test whether FleQ-mediated activation of *nrdR* depends on c-di-GMP, we performed fluorescent based reporter assay with and without exogenous c-di-GMP (0.1 mg/mL added at 4.5 h post-inoculation).

We compared the expression of wild-type and FleQ box mutated *PnrdR* and *PpelA* promoters using pETS130 based fusions (pETS130:P*nrdR*(PAO1 and *pelA* pETS130:P*pelA*(PAO1 and their respective mutated FleQ boxes (pETS130:P*nrdR*(PAO1) ΔFleQBox, pETS130:P*pelA*(PAO1) ΔFleQBox1, pETS130:P*pelA*(PAO1) ΔFleQBox2; see Materials and Methods). Fluorescence was recorded every 30 min post-treatment. We observed an increase in *nrdR* and *pelA* expression 2-3.5 h after c-di-GMP addition, particularly in FleQbox mutant constructs (Figure 5A-C). This suggests that FleQ-dependent *nrdR* activation is at least partially modulated by c-di-GMP, although other regulatory mechanisms may also contribute to this process.

**Fig 5.**
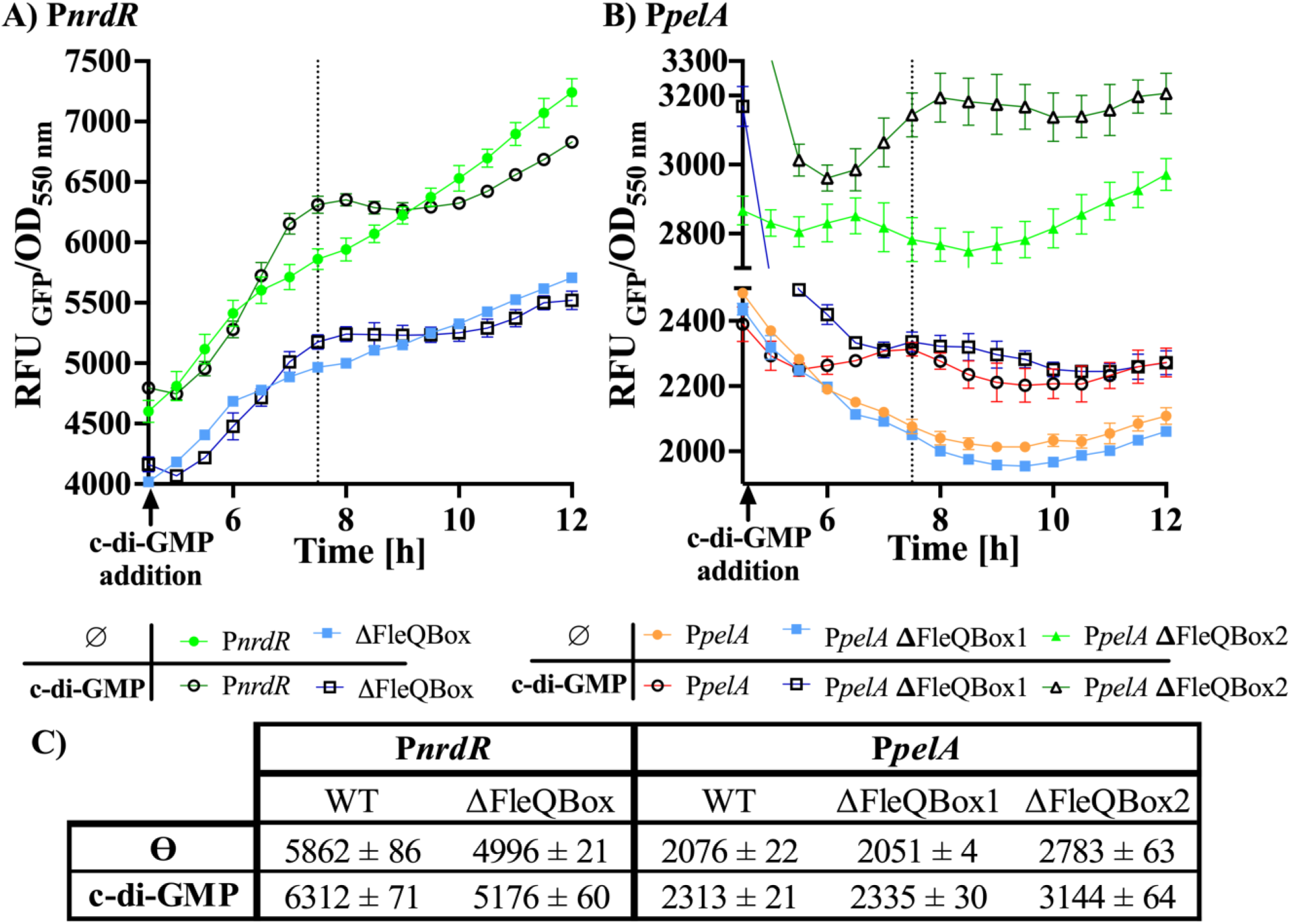
Impact of c-di-GMP on FleQ-dependent gene expression. A) *nrdR* promoter activity in the presence/absence of c-di-GMP (pETS130:P*nrdR*(PAO1), green and ΔFleQBox pETS130:P*nrdR*(PAO1) ΔFleQBox, blue). B) *pelA* promoter activity with or without 0.1 mg/ml c-di-GMP in wild type and FleQ box mutated constructs (pETS130:P*pelA*(PAO1), orange; P*pelA* ΔFleQBox1 [pETS130:P*pelA*(PAO1) ΔFleQBox1, blue and P*pelA* ΔFleQBox2 [pETS130:P*pelA*(PAO1) ΔFleQBox2, green). C) Fluorescence values (from the dotted line points) corresponding to 3 h post addition of 0.1 mg/mL c-di-GMP. Error bars indicate standard deviation (SD) from three independent replicates.

### NrdR regulation under biofilm-forming conditions

To investigate *nrdR* regulation during biofilm growth, we performed both static and continuous-flow biofilm formation experiments (see Materials and Methods), measuring total biofilm biomass and *nrdR* expression levels. In both models, we observed increased *nrdR* expression in FleQ box-mutated strains compared to the wild-type promoter, particularly at later time points that correspond to mature biofilms (Figures 6A and 7A–B). These results suggest that FleQ may also act as a repressor of *nrdR* during biofilm development.

**Fig 6.**
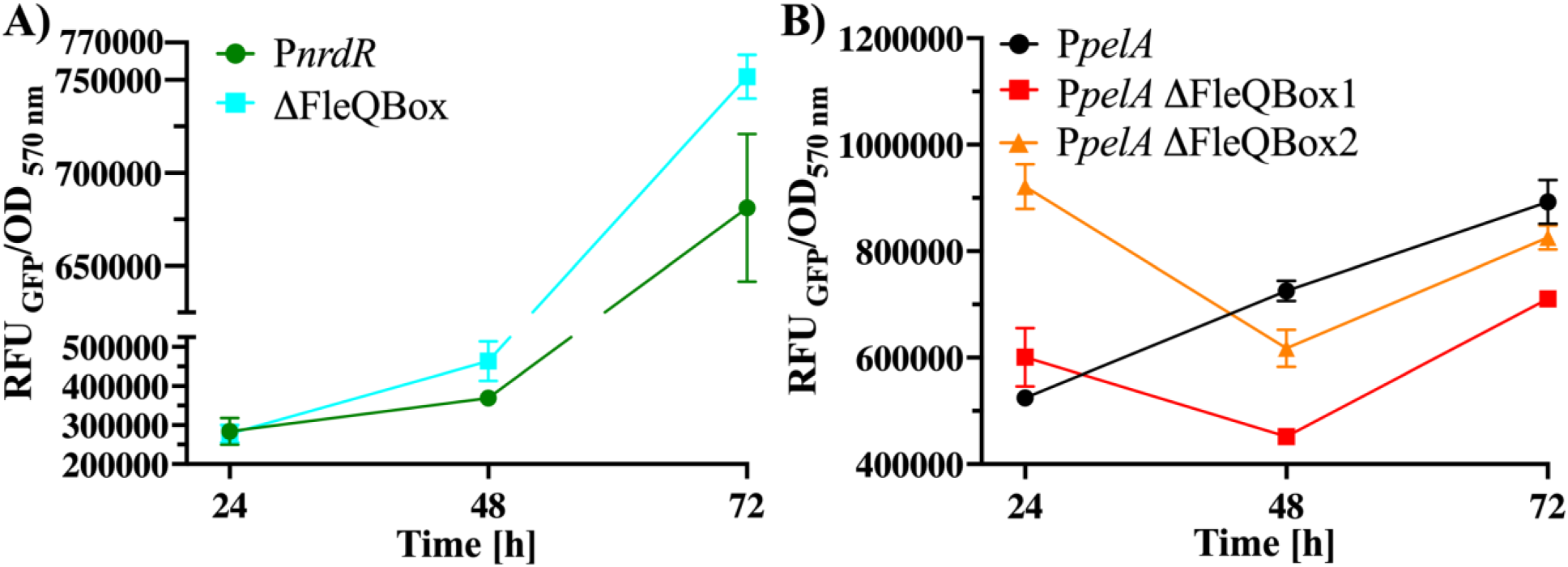
Static biofilm assays. GFP reporter measurements of (A) *nrdR* and (B) *pelA* promoter activity normalized to biofilm biomass (OD_570_) at 24, 48, and 72 h. Strains include wild-type and FleQ box-mutant constructs. Gene expression was measured using green fluorescent protein (GFP) as a reporter gene. *P. aeruginosa* PAO1 strains used: P*nrdR* [pETS130:P*nrdR*(PAO1), green], ΔFleQBox [pETS130:P*nrdR*(PAO1) ΔFleQBox, blue], P*pelA* promoter [pETS130:P*pelA*(PAO1), black], P*pelA* ΔFleQBox1 [pETS130:P*pelA*(PAO1) ΔFleQBox1, red] and P*pelA* ΔFleQBox2 [pETS130:P*pelA*(PAO1) ΔFleQBox2, orange].

**Fig 7.**
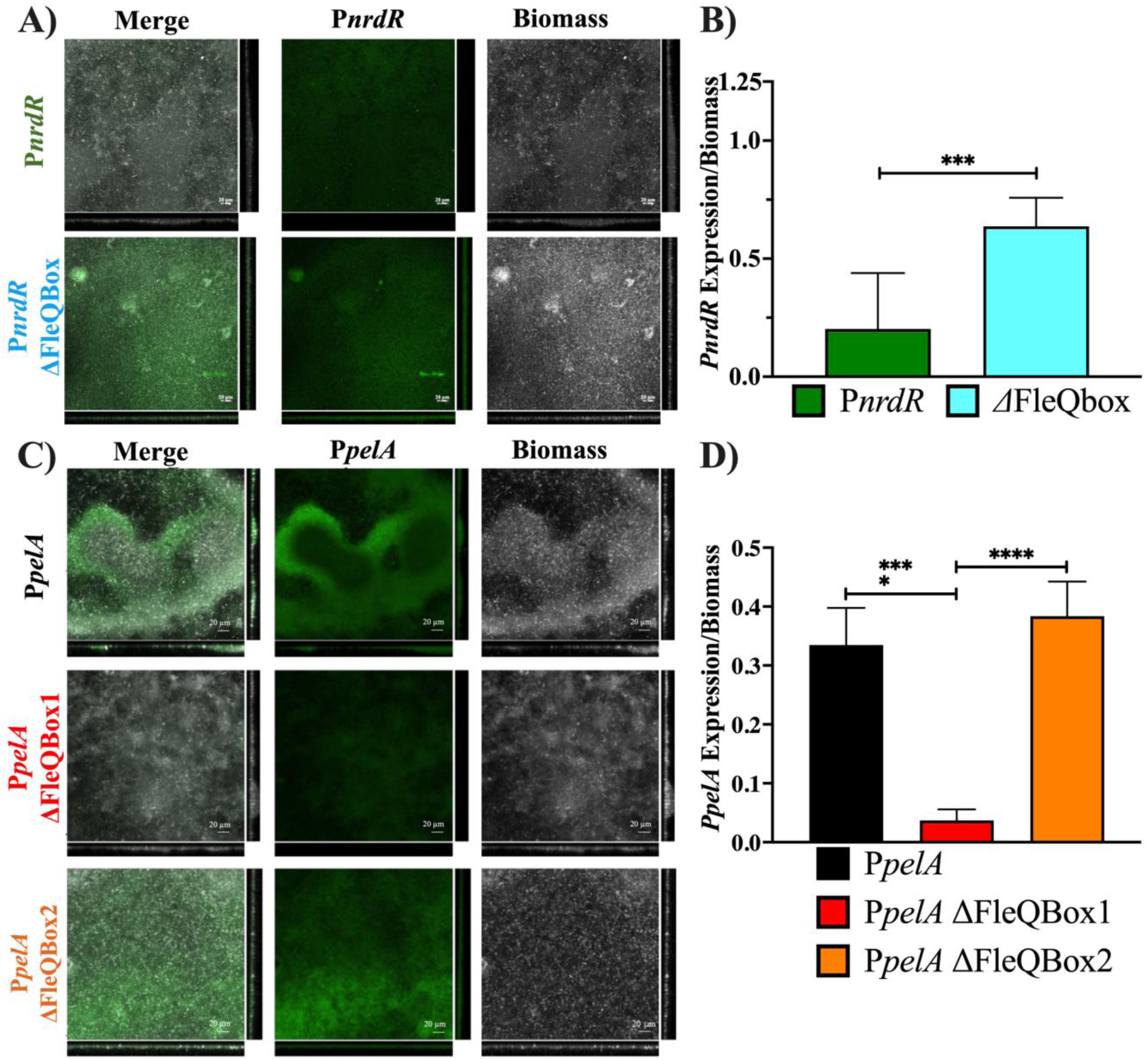
Continuous-flow biofilms expression studies of *nrdR* and *pelA*. A) CLSM images of continuous-flow *P. aeruginosa* PAO1 biofilms showing *nrdR* expression from the wild-type promoter (P*nrdR*) and the FleQ box mutated (P*nrdR* ΔFleQbox). B) Quantification of *nrdR* expression normalized to biomass in continuous biofilms. C) CLSM images of *P. aeruginosa* PAO1 biofilms expressing *pelA* from the wild-type promoter (P*pelA*), FleQ Box1-mutated promoter (P*pelA* ΔFleQBox1), and a FleQ Box2 mutated promoter (P*pelA* ΔFleQBox2). D) Quantification of *pelA* expression normalized to biomass in continuous biofilms. Biofilms were stained with SYTO60 (grey) and GFP is shown in green. Statistical analysis was performed using and unpaired Student’s *t*-test (***, *p* < 0.001; ****, *p* < 0.0001). *P. aeruginosa* PAO1 strains used: P*nrdR* [pETS130:P*nrdR*(PAO1), green], ΔFleQBox [pETS130:P*nrdR*(PAO1) ΔFleQBox, blue], P*pelA* promoter [pETS130:P*pelA*(PAO1), black], P*pelA* ΔFleQBox1 [pETS130:P*pelA*(PAO1) ΔFleQBox1, red] and P*pelA* ΔFleQBox2 [pETS130:P*pelA*(PAO1) ΔFleQBox2, orange] Scale bars correspond to 20 µm.

The *pelA* gene was used as a positive control to confirm FleQ function (Figures 6B, 7C-D). In the static biofilm experiment, the expression of the wild-type *pelA* promoter at 72 h was similar to that of the promoter carrying the FleQbox2 mutation (ΔFleQbox2), but higher than that of the promoter with FleQbox1 (Figure 6B). These observations were further corroborated by CLSM imaging (Figure 7C), which confirmed the reduced fluorescence intensity of the ΔFleQbox1 mutant (Figure 7D) compared to both the wild-type and ΔFleQbox2 constructs.

### NrdR expression under infection

*Galleria mellonella* is a widely used alternative model for studying bacterial infection due to its similarities to the mammalian innate immune system and practical advantages for *in vivo* assays. Previous studies have shown its suitability for monitoring gene expression during infection (26, 27). To investigate the regulation of *nrdR* in vivo, *P. aeruginosa* PAO1 strains carrying different promoter constructs fused to the *luxCDABE* operon (via the pETS220-BIATlux:PnrdR plasmids) were injected into *G. mellonella* larvae (see Materials and Methods). This allowed for real-time measurement of bioluminescence as a proxy for promoter activity at 15- and 18-hours post-infection.

As shown in Figure 8, the wild-type *PnrdR* construct displayed a 75.2-fold induction in expression between 15 and 18 hours. This induction was significantly reduced in the FleQ box- mutated construct (ΔFleQBox), which showed only a 44.2-fold increase. This indicates that FleQ positively regulates *nrdR* expression during infection. A mock construct containing a fragment of the *anr* gene (of similar size to the promoters used) was included as a negative control (pETS220- BIATlux:anr), confirming the specificity of the observed responses.

**Fig 8.**
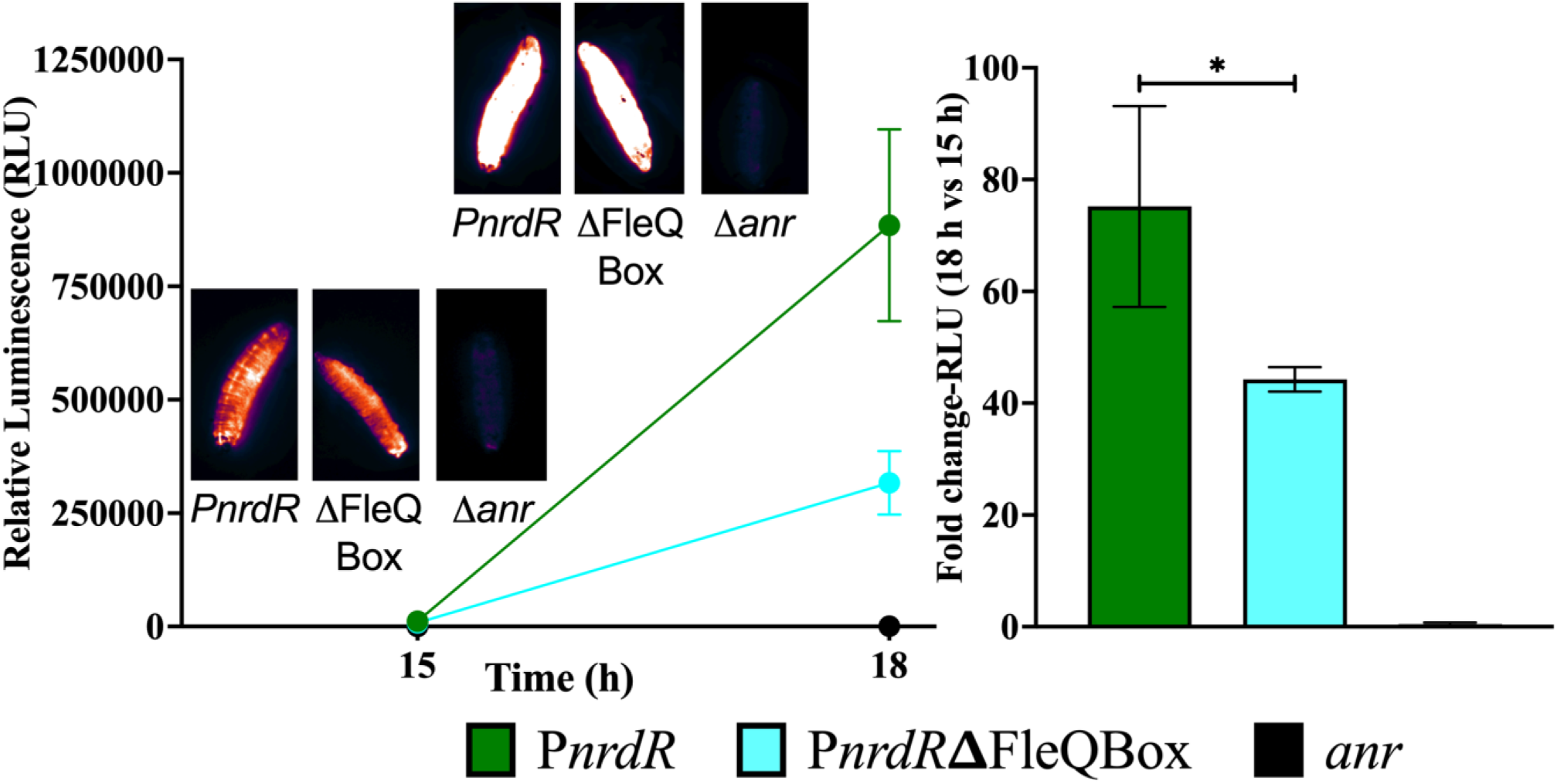
*nrdR* expression during *G. mellonella* larvae infection. A) Bioluminescent imaging of *G. mellonella* larvae infected with *P. aeruginosa* PAO1 strains expressing *luxCDABE* under the control of PnrdR or PnrdR ΔFleQBox, at 15 h and 18 h post-infection. Visualization was performed using the Gem LUT in ImageJ Fiji. B) Quantification of *nrdR* expression as fold-change between 15 h and 18 h post-infection. Strains used: P*nrdR* [pETS220-BIATlux:PnrdR, green], ΔFleQBox [pETS220-BIATlux:PnrdR ΔFleQBox, blue], and *anr* mock control [pETS220-BIATlux:anr, black]. Data represent the means of three independent experiments; error bars indicate the positive standard deviation. Statistical analysis was performed using an unpaired Student’s *t*-test (*, *p* < 0.05).

To further confirm the role of FleQ, a Δ*fleQ* mutant strain and its complemented counterpart were analyzed. The complemented strain restored *nrdR* expression to wild-type levels (see Supplementary Figure S6), supporting that the effects observed in the ΔFleQBox mutant were indeed due to the loss of FleQ-mediated regulation.

In addition to microplate-based luminescence quantification, a representative of individual larvae was imaged using the ImageQuant LAS 4000 system, and the bioluminescence pattern was consistent with the quantitative results (Figure 8A).

## Discussion

The medical relevance of *P. aeruginosa* has increased in recent years, particularly due to its association with nosocomial infections and chronic infections such as those in cystic fibrosis and other forms of bronchiectasis (1, 2). This pathogen’s remarkable environmental adaptability is largely driven by its genetic plasticity, allowing it to thrive under a wide range of conditions, including aerobic, anaerobic (32, 33), biofilm forming environments (34), and to infect diverse hosts including plants, animals, and humans (35).

This study focused on NrdR, a transcriptional regulator involved in controlling the expression of all three classes of ribonucleotide reductases (RNRs) in *P. aeruginosa* (11). Although *nrdR* was first identified in *S. coelicolor* (13, 15) and subsequently in *E. coli* (36), mycobacteria (37) and *P. aeruginosa* (11), the regulation of this gene remains incompletely understood. Previous work from our group identified NarL as a transcriptional activator of *nrdR* under anaerobic conditions (11). Here, we aimed to expand this knowledge by identifying additional transcriptional factors (TFs) that modulate *nrdR* expression under various conditions: aerobic, anaerobic, biofilm growth, and during *G. mellonella* infection.

First, we mapped the transcription start site (TSS) of *nrdR* using 5’ RACE, locating it -28 bp upstream of the start codon (Figure 1). This result is consistent with previous transcriptomic studies under both aerobic and anaerobic conditions (28). Bioinformatic analysis identified several putative TF binding sites with the *nrdR* promoter region (Figure 1). Among these, quorum sensing (QS) boxes were excluded due to the lack of expression changes in QS mutant strains (Supplementary Figure S2). Two NarL binding sites had already been experimentally confirmed by our group under anaerobic conditions (11).

Importantly, we identified a FleQ binding site located approximately 224 bp upstream of the TSS. Although this is a relatively distant, other FleQ-regulated genes (e.g. *fleSR*) have been shown to be regulated via DNA looping (31, 38). Similarly, FleQ binding sites at -294 and -263 bp upstream of PA2440 have been reported (39), supporting the feasibility of FleQ-mediated regulation of *nrdR*.

To validate these bioinformatic predictions, we attempted electrophoretic mobility shift assays (EMSAs) but could not obtain soluble FleQ despite multiple solubilization strategies. As an alternative, we used a pull-down assay with tandem repeats of the *nrdR* promoter, crude extract from *P. aeruginosa* PAO1, and magnetic beads (Supplementary Figure S4) to directly isolate the proteins that bind to the *nrdR* promoter region. Proteomic analysis via LC-MS confirmed the direct interaction of FleQ and NarL with the *nrdR* promoter and also identified other potential regulatory proteins that need further studies. These findings offer a valuable starting point for future studies on the transcriptional regulation of *nrdR*.

During the exponential growth, RNRs are essential for deoxyribonucleotide synthesis, which supports DNA replication and repair. Tight regulation of this process is therefore crucial. NrdR has previously been established as a key repressor of RNR genes (11, 17, 18, 36, 40). Here, we show that FleQ acts as an activator of *nrdR* under planktonic conditions, both aerobically and anaerobically (Figure 3-4). The Δ*fleQ* mutant exhibited slower anaerobic growth, consistent with FleQ’s involvement in regulating genes required for anaerobic respiration, similar to its role in denitrification in *Pseudomonas fluorescens* (41). RT-qPCR revealed significantly reduced *nrdR* expression in Δ*fleQ* cells (Figure 3B).

To further examine FleQ regulation, we used promoter-probe assays with wild-type and mutant *nrdR* promoters (ΔFleQBox). These experiments, complemented by plasmid-based FleQ expression, confirmed that FleQ functions as a positive regulator of *nrdR* transcription (Figure 4). The effect was abolished in the FleQ box mutant and restored upon complementation, providing strong support for direct regulation. *pelA* served as a control for FleQ’s transcriptional regulation (29, 30).

Since FleQ activity is modulated by intracellular cyclic-di-GMP (c-di-GMP) levels (29–31), which is involved in regulating virulence and biofilm formation in bacteria (42), we next tested whether this second messenger affects *nrdR* expression. Under high c-di-GMP conditions (biofilm growth) (30, 42), we observed increased *nrdR* expression (Figure 5), although this induction was less pronounced than for the known FleQ target *pelA* (29). This difference may be due to the presence of additional FleQ binding sites in the *nrdR* promoter, a hypothesis supported by the partial regulatory response observed. FleQ is known to switch between activator and repressor functions depending on intracellular c-di- GMP concentrations (29–31, 38). In biofilm conditions, where c-di-GMP is elevated, FleQ acts as a repressor of *nrdR,* exhibiting an opposite response observed during planktonic growth (Figure 4, 6-7). This was evident from both static and flow biofilm models (Figures 6–7). In static biofilms, *nrdR* expression increased progressively over three days, with even higher expression in FleQ box mutants (Figure 6). A Δ*fleQ* strain was not possible to use in these studies because it cannot form a mature biofilm (30). We used the P*pelA* promoter as a positive control for our biofilm experiments, focusing on the second FleQ box due to its stronger gene expression response (29). The continuous flow-cell biofilm model corroborated these results (Figure 7), aligning with known characteristics of chronic biofilm infections (43).

We then evaluated *nrdR* expression *in vivo* using the *G. mellonella* infection model (26, 44–46). While this system mimics acute infections and supports analysis of RNR expression (27), it is less suitable for studying biofilms due to immune-mediated bacterial clearance and the short lifespan of infected larvae that don’t allow for mature biofilm development (47). Thus, *nrdR* expression in *G. mellonella* reflects planktonic growth, rather than biofilm conditions. Using P*nrdR*-lux constructs with wild-type and mutant FleQ boxes, we demonstrated that FleQ activates *nrdR in vivo* at 18 hours post-infection (Figure 8), consistent with the results of planktonic growth (Figure 4). Expression was significantly reduced in the FleQ box mutant and restored by FleQ complementation (Figures 8). Interestingly, the *ΔfleQ* strain showed no difference in larval survival compared to WT (Supplementary Figure S6), but *nrdR* expression was clearly affected, confirming FleQ’s role as an activator during infection.

Figure 9 provides a graphical summary of the regulatory pathway governing *nrdR* expression in *P. aeruginosa* under different conditions. FleQ modulates *nrdR* in response to environmental cues, particularly through c-di-GMP levels, acting as an activator in planktonic and infection conditions (related to acute infections) and as a repressor in mature biofilms. This dual function is consistent with FleQ’s role as master regulator of the transition from motile to sessile states and modulated by intracellular c-di-GMP levels (30, 31, 38, 39, 48), which are synthesized by diguanylate cyclases (DGCs) and degraded by phosphodiesterases (PDEs) (Figure 9). Its activity is further influenced by others regulators, such as FleR (via the FleS/FleR two-component system) (49) and the Vfr repressor (50) (Figure 9).

**Fig 9.**
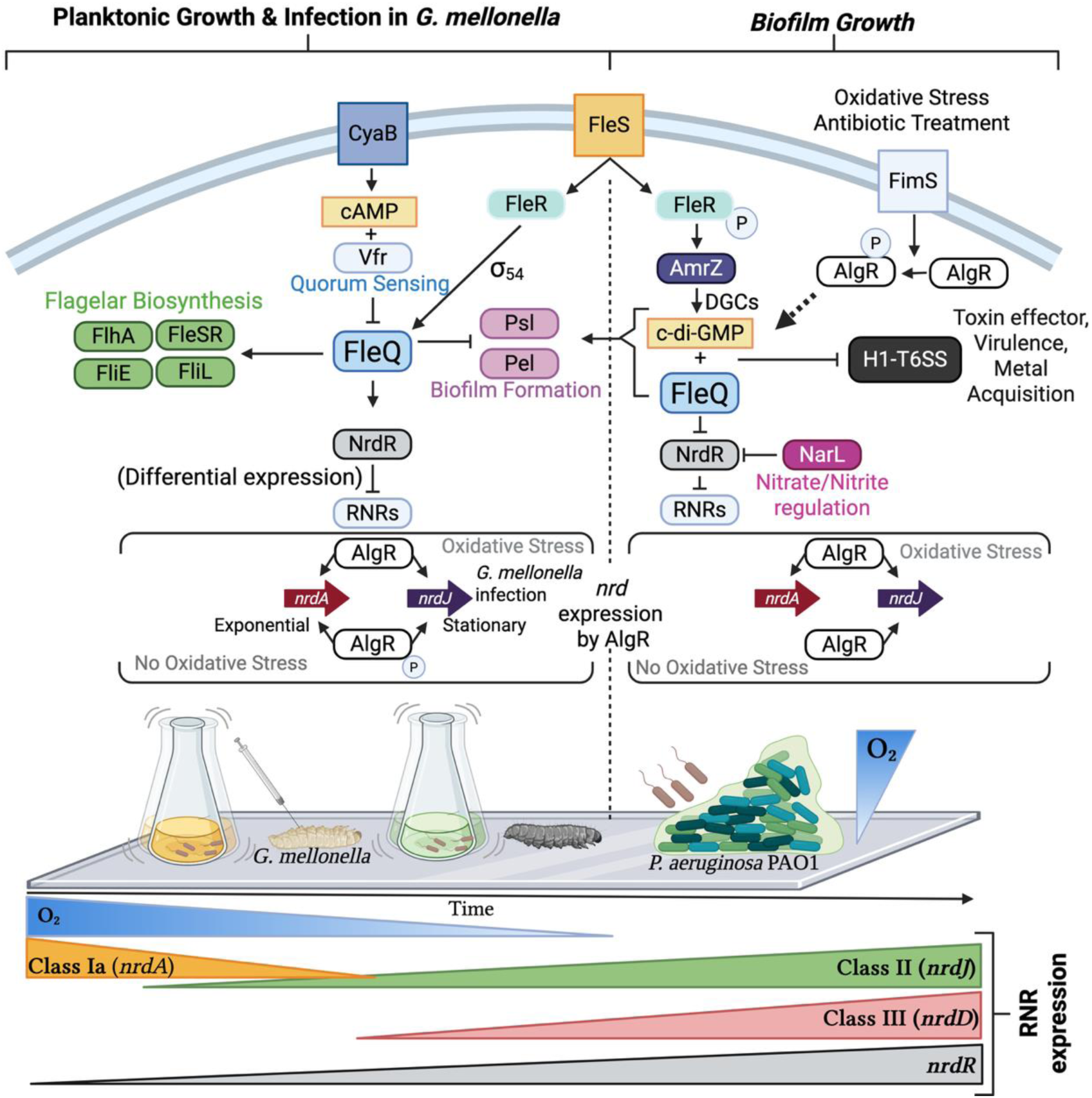
Graphical representation of the *nrdR*/FleQ regulatory pathway in *P. aeruginosa* PAO1 and *G. mellonella*. The regulatory pathway is depicted under planktonic growth conditions and during *G. mellonella* infection. FleQ functions as an activator of both *nrdR* and flagellar biosynthesis genes (e.g. *fleSR, flhA, fliE, fliL, etc.*), while simultaneously repressing biofilm formation genes (*psl, pel,* etc.). FleQ itself is regulated by quorum-sensing (QS) proteins such as Vfr or FleR. During biofilm formation, FleQ and NarL act as repressors of *nrdR*. Under these conditions, FleQ regulation is influenced by the intracellular concentration of c-di-GMP, which is elevated due to the upregulation of diguanylate cyclase (DGC) by AmrZ and oxidative stress signal (FimS-AlgR). The regulation of *nrdA* and *nrdJ* by AlgR is highlighted in boxed sections. At the bottom of the image, the expression levels of ribonucleotide reductase (RNRs) and the fold-change in *nrdR* gene expression over time are shown. The image was created using BioRender.com.

Given the essential nature of *nrdR* (11) and FleQ’s central role in the planktonic-to-biofilm transition (38, 48), it is not surprising that their regulatory interplay is tightly coordinated.

FleQ, in conjunction with c-di-GMP, modulates *nrdR* transcriptional, especially during mature biofilm formation (Figures 6-7) and the stationary phase (Figure 4) when *nrdR* is essential for repressing RNRs (11, 17, 40). Under low c-di-GMP conditions (30, 31, 38, 39, 48), FleQ acts as a master regulator in *P. aeruginosa*, activating flagellar genes (*flhA*, *flhE*, *fliL*, and *fleSR*) (51) during planktonic conditions while repressing biofilm-associated genes (*pel, psl*) (Figure 9).

*P. aeruginosa* requires both dispersal mechanisms (via flagellar genes) and tight regulation of RNRs, controlled by factors such as NrdR, AlgR among others, as nucleotide synthesis is crucial for DNA repair and de novo synthesis (6). During exponential growth, RNR class Ia is highly expressed, while classes II and III are repressed (Figure 9). Under these conditions, FleQ promotes the expression of both *nrdR* and *pelA,* not only during exponential growth but also during infection in *G. mellonella*, resembling an acute infection (Figures 3, 4, 5, 8). In contrast, during biofilm formation, FleQ repressed *nrdR* (Figures 6-7), likely to reduce DNA synthesis and metabolic activity, hallmarks of biofilm associated persistence (43, 52).

In biofilms, cells prioritize matrix production, driven by increased c-di-GMP levels through the activation of two-component systems such as FleS/FleR (49) and FimS/AlgR (27, 53) (Figure 9). This second messenger is crucial for FleQ-mediated biofilm synthesis via the *psl* and *pel* (38) gene clusters (Figure 9). Chronic infections characterized by mature biofilms results in the repression of toxin secretion, virulence factors, and metal acquisition (54). FleQ, in coordination with c-di-GMP, contributes to this repression by downregulating the H1-T6SS (49, 51) (Figure 9).

Previous work from our group demonstrated that NarL also acts as a positive regulator of *nrdR* under anaerobic and biofilm conditions (11). This highlights the complexity of *nrdR* regulation, with multiple inputs modulating expression in response to environmental conditions. Notably, FleQ’s regulatory logic at the *nrdR* promoter resembles that seen for *pelA* (Figures 6-7) and *fleSR*, where binding sites mediate both activation and repression depending on c-di-GMP levels (38, 48).

In conclusion, this study significantly advances our understanding of *nrdR* regulation in *P. aeruginosa* (Figure 9). We have identified new putative transcriptional regulators via pull-down assays and demonstrated that FleQ is a key modulator of *nrdR* expression, acting as an activator in planktonic growth and acute infection, and a repressor in biofilm-promoting conditions. This regulation is mainly independent of oxygen availability, consistent with *nrdR*’s essential role under diverse environmental conditions (11). Our findings offer new insights into the transcriptional network controlling RNR activity and establish FleQ as a critical link between environmental adaptation and nucleotide metabolism in *P. aeruginosa*.

## Methods

### Bacterial strains and growth conditions

The bacterial strains used in this study are listed in Supplementary Table S1. Routine cultures were grown in Luria-Bertani (LB) broth at 37 °C. *Escherichia coli* BL21(DE3) was used for protein overproduction, while *E. coli* DH5α was used for plasmid construction. Anaerobic growth of *P. aeruginosa* was performed in screwcap Hungate tubes filled to the brim with LB supplemented with 1% (w/v) KNO_3_ (LBN) and saturated with N_2_. Antibiotics were added at the following concentrations when required: for *E. coli*, 10 µg ml^-1^ gentamicin, 17 µg ml^-1^ chloramphenicol, 50 µg ml^-1^ kanamycin, 50 µg ml^-1^ ampicillin; for *P. aeruginosa*, 100 µg ml^-1^ gentamicin, 300 µg ml^-1^ carbenicillin. Where indicated, 0.1 mg/mL of cyclic di-GMP (Sigma) was added.

### DNA manipulation and plasmid construction

All plasmids and primers used in this study are listed in Supplementary Table S1 and S2, respectively. Genomic DNA (gDNA) from *P. aeruginosa* PAO1 was amplified using Phusion High- Fidelity DNA Polymerase (ThermoFisher Scientific) according to the manufacturer’s instructions. Plasmid extractions were performed using the GeneJET Plasmid MiniPrep Kit (ThermoFisher Scientific). Restriction digestion and ligation were carried out using enzymes from Thermo Fisher Scientific and T4 DNA ligase (Invitrogen), respectively. PCR and digested products were purified using the GeneJET Gel Purification Kit (ThermoFisher Scientific) according to the manufacturer’s instructions.

PCR products were initially cloned into pJET1.2 plasmid (ThermoFisher Scientific) and subsequently subcloned into pETS130, pUCP20T, or pETS220-BIATlux using the following enzyme pairs: *Bam*HI/*Cla*I, *Xba*I, or *Sac*I/*Eco*RI, respectively. Correct constructs were verified by colony PCR using DreamTaq Master Mix (ThermoFisher Scientific) with primers 1/2 for pETS130, pUCP20T and pETS220-BIATlux, and primers 3/5 for pJET1.2 constructs. *P. aeruginosa* was transformed by electroporation, as described previously (11).

### Bioinformatic identification of transcriptional factors

The 355 pb region upstream of the *nrdR* start codon (PA4057), hereafter referred to as P*nrdR*, was from the *Pseudomonas* Genome DB (ID: 110934) (www.pseudomonas.com) (see Figure 1). Putative transcription factors (TF) binding sites were predicted using the PRODORIC2 Virtual Footprinting tool (19) with all *P. aeruginosa* weight matrices (PWMs). Hits were considered positive if the score exceeded 5 and the binding site was on the sense strand (see Figure 1C).

### Site-directed mutagenesis of Transcriptional factors binding sites

Putative FleQ and NarL binding sites identified within *PnrdR* were site-directed mutated by overlap extension PCR (20) using primers 6 to 11. Mutagenesis of FleQ binding sites with P*pelA*, used as a positive control, was performed similarly using primers 12 to 16, as previously described (11).

### FleQ overproduction

FleQ was overproduced in *E. coli* BL21 (DE3) using pET28a plasmid (Novagen). The *fleQ* coding region from *P. aeruginosa* PAO1 was amplified with primers 17/18 and cloned into pET28a to generate pET28a:*fleQ.* Cultures were inoculated at OD_550_ = 0.05 and grown to OD_550_ = 0.5 before induction with 0.1–1 mM IPTG, with or without 0.1 mg of c-di-GMP. Cultures were incubated for 3–4 h at 37 °C or overnight at 15 °C. Crude extracts were prepared using BugBuster reagent (Sigma) and analyzed by 7.5% SDS-PAGE (Mini-PROTEAN® TGX™, Bio-Rad) with Coomassie staining.

### RNA extraction, RT-PCR, and transcriptional start site determination

Strains were cultured to OD_₅₅₀_ = 0.4–0.6 (exponential phase) and 2.2–2.5 (stationary phase), harvested, and stabilized in 500 μL RNAlater (Thermo Fisher Scientific). RNA was extracted using the RNeasy Mini Kit (Qiagen) and treated with DNase I (ThermoFisher Scientific) to remove contaminant DNA. cDNA was synthesized from 1 μg of total RNA using iScript (Bio-Rad). RT-PCR was performed using SYBR Green Master Mix (ThermoFisher) with 200 nM primer pairs 19/20, 21/22, and 23/24, and analyzed on an ABI Step One Plus system (Applied Biosystems). Expression was normalized to *gapA* (21).

Transcriptional start site (TSS) mapping of the *nrdR* gene was performed by 5’-RACE using RNA from aerobic and anaerobic cultures (OD_600_ of 0.5). cDNA was polyadenylated using Terminal Deoxynucleotidyl Transferase (ThermoFisher Scientific), and two rounds of PCR were performed using primer polyT (primer 29) and primer 25 (PCR1) and 2polyT (primer 30) and primer 6 (PCR2). Amplicons were cloned into pJET1.2, transformed into *E. coli* DH5α. PCR screened, and sequenced using primers 3/5 and the orientation of the cloned fragment was determined using primer 26.

### Transcriptional expression during planktonic and biofilm growth

Wild-type and mutant versions of P*nrdR* and P*pelA* were cloned into the GFP-reporter plasmid pETS130^23^, generating pETS130:P*nrdR*(PAO1), pETS130:P*nrdR*(PAO1) ΔFleQBox, pETS130:P*nrdR*(PAO1) ΔNarLBox1&2. We did the same for P*pelA* promoters, generating the following vectors: pETS130:P*pelA*(PAO1), pETS130:P*pelA*(PAO1) ΔFleQBox1, pETS130:P*pelA*(PAO1) ΔFleQBox2. GFP fluorescence (λ_ex 485 nm / λ_em 535 nm) was measured during cells grown in exponential (OD_550_ = 0.45–0.55) and stationary (OD_550_ = 2.5–4.0) phases and normalized to OD_550_ (22).

For static biofilms, 3-day cultures were grown in 96-well microtiter plates (Corning, Inc. Costar). Fluorescence was normalized to biomass quantified by crystal violet staining, with OD_570_ used as the readout according to previous work (22).

For continuous biofilm, cultures were adjusted to an OD_550_ of 0.3 and inoculated into three- channel flow cells and incubated for 4 h. LB medium supplemented with 0.2% glucose was flowed at a constant flow rate of 42 μL/min using an Ismatec ISM 943 peristaltic pump (Ismatec, Wertheim, Germany) (23). After 72 h, biofilms were stained with 10 µM SYTO60 for 30 min (Thermo Fisher Scientific, MA, USA) and imaged using a Zeiss LSM 800 confocal laser scanning microscope (CSLM) with excitation wavelengths of 488 and 640 nm. Microscope images were processed using ImageJ and COMSTAT 2 software was used to quantify biofilm biomass (24).

To complement the *P. aeruginosa* PAO1 *ΔfleQ* mutant, the *fleQ* coding region with its native promoter was obtained by PCR with primers 27/28 and digestion with *Xba*I and *Bam*HI, cloned into pUCP20T to generate the pUCP20T:P*fleQ*:*fleQ* plasmid.

### Pseudomonas aeruginosa PnrdR promoter pull-down assay

A synthetic *PnrdR* sequence (triplicated in tandem) was cloned into pJET1.2b using GeneArt platform (ThermoFisher), generating pMX-P*nrdR*(PAO1)x3. A 5’ biotin-labeled PCR product was amplified using primers 4/5 and used in pull-down assays with streptavidin-magnetic beads (M-280 Dynabeads, Thermo Fisher) as previously described (25) (See Supplementary Figure S1).

Protein extracts from *P. aeruginosa* cultures (OD_550_ = 0.55 or >2.5) were incubated with beads, and bound proteins were resolved by 12% SDS-PAGE (Mini-PROTEAN® TGX™ Precast Gel- BioRad) and visualized via Coomassie staining. Digital images of the SDS-Page gel were captured with an ImageQuantTM LAS 4000 mini-imager (GE Healthcare, Chicago, IL, USA) with a 1/30 second exposure time. Bands were excised, digested with trypsin, and analyzed by liquid chromatography-mass spectrometry (LC-MS) using tandem a 60-min gradient on an Orbitrap Eclipse instrument at the Proteomics and Metabolomics Service at the Germans Trias i Pujol Research Institute (IGTP). As a quality control, BSA was digested in parallel, run between samples to avoid carryover, and assess the instrument performance.

Peptides were identified using Proteome Discoverer v2.5 and Mascot v2.6 with the *P. aeruginosa* PAO1 proteome (UP000002438), filtered for false Discovery rate (FDR) < 1%. Results are available in the Supplementary Table III.

### *Galleria mellonella* maintenance and injection

*G. mellonella* larvae were reared at 34°C in darkness on an artificial diet (15% corn flour, 15% wheat flour, 15% infant cereal, 11% powdered milk, 6% brewer’s yeast, 25% honey, and 13% glycerol) (26, 27). Overnight bacterial cultures were washed three times and resuspended in PBS (Fisher Scientific, Madrid, Spain), and OD_590_ was adjusted to 1. 10-fold serial dilutions were prepared and injected (5–20 CFUs per larva) into the top right proleg using a 26-gauge microsyringe (Hamilton, Reno, NV, USA). Groups of 6 larvae were infected per condition and incubated at 37 ^◦^C.

### P*nrdR*-Lux transcriptional fusions and bioluminescence measurement in *G. mellonella*

Transcriptional fusions between P*nrdR* (or its mutated variants) and *lux* reporter gene were generated in pETS220-BIATlux plasmid, yielding pETS220-BIATlux:P*nrdR* and pETS220- BIATlux:P*nrdR* ΔFleQBox. Larvae were anaesthetized on ice for 10 minutes before relative bioluminescence measurements at 15 and 18 h post-infection using a 6-well microtiter plate (Caplugs Evergreen, Buffalo, NY, USA) and a 200 Pro Fluorescence Microplate Reader (integration time 1000 ms) (Tecan, Switzerland). Bioluminescent images were captured with an ImageQuantTM LAS 4000 mini-imager (GE Healthcare, Chicago, IL, USA), with a 30-second exposure time and processed using FIJI/ImageJ software (Version 1.52p, National Institutes of Health, Bethesda, MD, USA) (26, 27).

## Acknowledgements

This study was partially supported by grants PID2021-125801OB-100, PLEC2022-009356 and PDC2022-133577-I00 funded by MCIN/AEI/10.13039/501100011033 and “ERDF A way of making Europe”, the CERCA programme and AGAUR-Generalitat de Catalunya (2021SGR01545), the European Regional Development Fund (FEDER) and the Catalan Cystic Fibrosis association. E.T. is a researcher of the ICREA Academia 2025 program. D.M. and J.A. thanks Generalitat de Catalunya for its financial support through the FI program (2020-FI-B-00175 and 2021-FI-B-00118).

The funders had no role in study design, data collection and interpretation.

## Author Contribution

The manuscript was written through the contributions of all authors. D.M, A.R, L.P, J.M.H, and J.A performed all the experiments and wrote the manuscript. E.T. conceived and supervised the research, revised the experimental data, and wrote the manuscript. E.T. directed and administered the project, and acquired the funding. All authors have approved the final version of the manuscript.

